# Differential Boric Acid and Water Transport in Type I and Type II Pores of Arabidopsis Nodulin 26-Intrinsic Proteins

**DOI:** 10.1101/2022.10.05.510970

**Authors:** Zachary G. Beamer, Pratyush Routray, Rupesh Agrawal, Tian Li, Katey M. Gibson, Katherine E. Ostrouchov, Jeremy C. Smith, Daniel M. Roberts

## Abstract

Nodulin-26 intrinsic proteins (NIPs) are plant-specific multifunctional aquaporin-like channels that are phylogenetically and structurally segregated into three subfamilies: NIP I, II, and III. Each subfamily has a characteristic selectivity filter sequence (the “aromatic-arginine” region, or ar/R) that controls substrate transport specificity based on steric constraints, hydrophobicity, and the spatial orientation of hydrogen bonding moieties. All three NIP subfamilies transport metalloid hydroxides, both beneficial as well as toxic, but with different selectivities. Here we investigated the B, As, and water selectivity of representative *Arabidopsis thaliana* NIP I and II proteins as well as their ar/R mutants in transport assays as well as through B complementation analysis in the B sensitive *nip5;1* mutant background. All NIP proteins, and their ar/R mutants, showed equal permeability to arsenite, but showed differences in boric acid and aquaporin activities that was linked to the amino acid at the helix 2 (H2) position of the ar/R filter (Ala for NIP II and Trp for NIP I). The presence of an alanine at this position in NIP II proteins enhances boric acid permeability and drastically reduces the aquaporin/water permeability of the channel. A NIP II structural model generated from the AlphaFold2 resource and evaluated by MD simulation shows that the alanine results in a wider ar/R pore that accommodates the trigonal boric acid molecule and may allow gating of the pore in a manner that affects water permeability. In contrast, NIP I proteins adopt a more classical aquaporin/glyceroporin arrangement in the ar/R that allows metalloid permeability, although with greater selectivity, as well as permeation by water.

## 1. Introduction

The aquaporin superfamily is an ancient family of structurally conserved integral membrane protein channels that facilitate the transport of water and uncharged solutes across cellular membranes. Each subunit of these homotetrameric proteins has a common fold that consists of six transmembrane α-helices with two conserved helical loop regions (NPA boxes) that fold back into the membrane forming a pore through the center of each subunit [reviewed in (1–3)]. Transport selectivity is determined by a selectivity filter termed the “aromatic/arginine” (ar/R) region (4) that is formed by the confluence of four amino acid residues: one each from α-helices 2 (H2) and 5 (H5), as well as two from the second NPA box (LE1 and a conserved arginine). The ar/R amino acids determine transport selectivity based on size, hydrophobicity and hydrogen bonding with transported substrates (5–8).

The evolution of land plants led to an expansion and diversification of the aquaporin gene family including the emergence of a land plant-specific subfamily termed the “Nodulin 26-like intrinsic proteins” (NIPs). NIPs are phylogenetically and structurally organized into three broad families, NIP I, II and III, that have distinct transport selectivities (9–14). NIP III proteins are widely distributed among the Graminaceae, and biochemical and genetic evidence show that they principally function as silicic acid channels (Si[OH]_4_) that promote optimal growth and development, as well as resistance to abiotic and biotic stress (15–18). NIP II proteins are boric acid channels that facilitate the uptake and distribution of this micronutrient to developing tissues under B limiting conditions (19–21). The biological and transport roles of NIP I proteins are less precisely defined, although myriad functions are postulated, ranging from metalloid permeability (22–26), to aquaporin and ammoniaporin transport during symbiosis (27–29), to lactic acid efflux during anaerobic stress (30,31). Each NIP pore subfamily possesses signature ar/R amino acid compositions that are postulated to determine their disparate substrate specificities and biological functions [reviewed in (14)]. Insight into the unique pore features and potential regulatory features of the NIP family has recently emerged with the solution of two atomic resolution crystal structures of an open and closed conformation of the OsNIP2;1/Lsi1 silicic acid channel (a NIP III protein) from rice (32,33). Structural information for NIP I and II proteins is not yet available.

The objective of this study was to examine the metalloid and aquaporin selectivity, and the function of signature ar/R residues, of NIP Type I and Type II proteins from Arabidopsis by evaluating the transport of three key NIP solutes – water, boric acid and arsenite, and by complementation analyses *in planta* utilizing the B-sensitive *nip5;1* mutant. In addition, to interpret the functional data from a structural perspective, NIP I and II models were generated by using the Alphafold2 method followed by MD simulation refinement and comparison to high resolution open and closed structures of the Lsi1 NIP III protein.

## 2. Results

### 2.1 The NIP fold and selectivity filter deviate from water-selective aquaporins

Insight into the NIP structure and how it has diverged from classical water specific aquaporins has come from the recent solution of the rice silicic acid transporter Lsi1 structure at atomic resolution (32,33). Lsi1 is a NIP III protein that contains the signature sequence of silicic acid channels with a wide and hydrophilic ar/R (Fig 1) with a consensus sequence of G-S-G-R [H2, H5, LE_1_, R] (Fig. S1). This property leads to three essential features that distinguish NIP III from other aquaporin and aquaporin-like proteins. First, the invariant serine at H5 is unique to NIP III proteins and provides a hydrogen bond donor to transported substrate. Second, the relatively open nature of the NIP III ar/R allows the insertion of a fifth residue from helix 1, a conserved threonine, into the selectivity filter, providing an additional ligand for transported substrates and bound water (Fig. 1B). Third, unlike water-specific aquaporins, which narrow to the diameter of a single water molecule at the ar/R allowing only single file water transport, Lsi1 is highly hydrated (Fig. S1) with three waters within the ar/R selectivity filter (blue in Fig. 1B), as well as two additional tightly-bound waters (green in Fig. 1B). These features are postulated to account for transport specificity by providing steric accommodation and multiple hydrogen bond contacts with Si(OH)_4_ as it traverses the pore (32,33).

**Figure 1.**
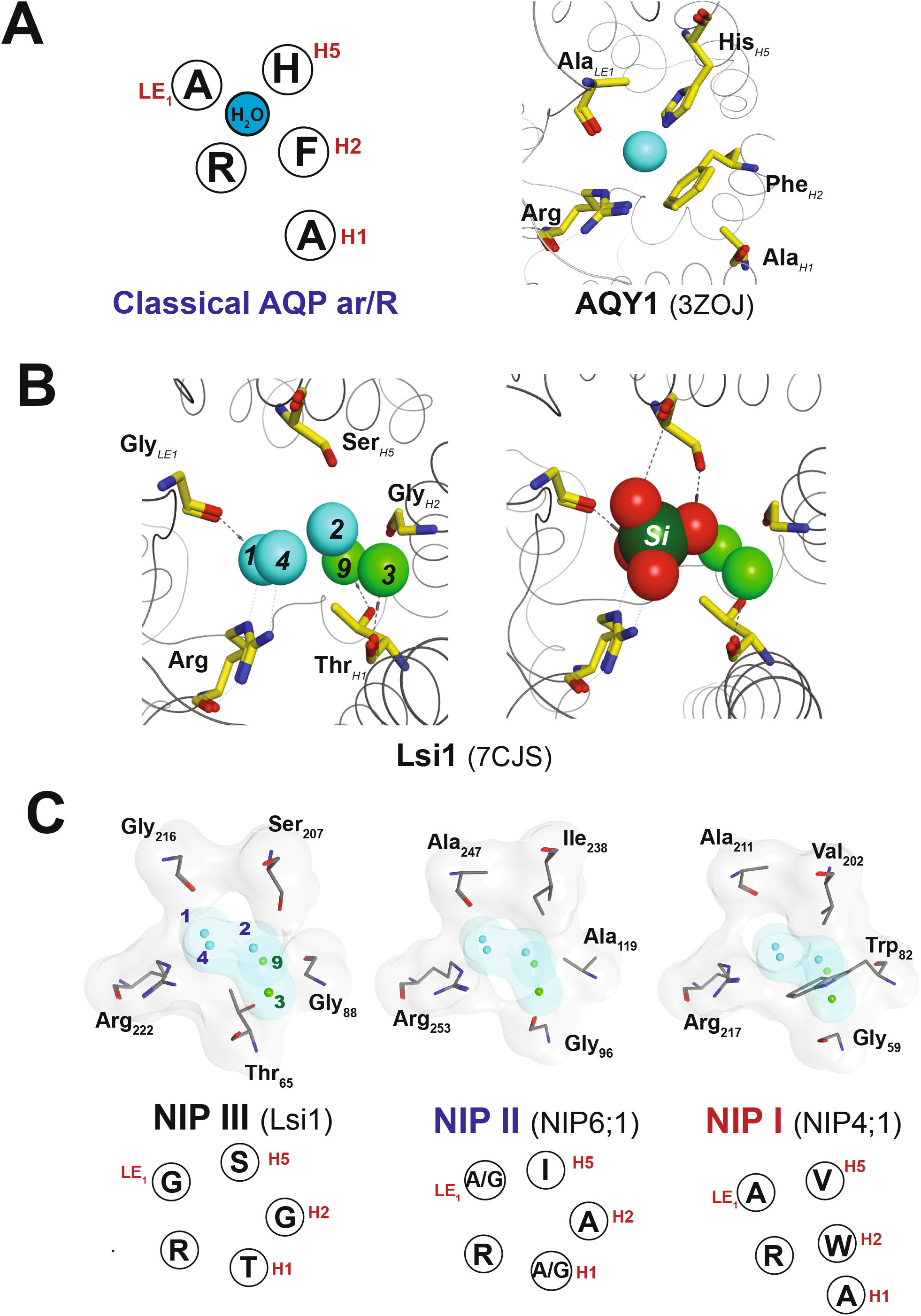
Representative models of the selectivity filter regions of NIP pore subtypes. **A.** *Left*, diagram of the ar/R selectivity filter for classical water specific aquaporins viewed down the pore axis from the extracellular vestibule with the ar/R nomenclature of Wallace and Roberts, (46) (H1: helix 1, H2: helix 2, H5, helix 5, LE_1_: E loop residue, R: conserved arginine). *Right*, **Aqy1**, the selectivity filter of yeast aquaporin Aqy1. **B.** *Left*, **Lsi1**, the rice NIP III silicic acid permease viewed from the same orientation with waters shown as aqua (transported waters) or green (tightly bound waters) spheres. The waters are numbered based on the nomenclature of Saitoh et al. (32). *Right*, **Lsi1** with silicic acid bound at the ar/R selectivity filter generated by QM/MM (32). **C.** The ar/R selectivity filters of the Lsi1 (NIP III) and the NIP6;1 (NIP II) and NIP4;1 (NIP I) homology models shown with the predicted positions of the ar/R bound waters based on the Lsi1 structure (32). The diagram below each structure shows the consensus ar/R sequence for the indicated NIP I, II, and III pore subtype based on phylogenic analyses across plant species (Fig. S1) (14).

To investigate the comparative pore properties of NIP I and NIP II proteins, in light of the features of the high resolution Lsi1 structure, structural models were generated from representative Arabidopsis NIP I and II proteins (Fig. S1). Initial structures were obtained using Alphafold2 (34), and were further refined by molecular dynamics simulation to generate a hierarchal cluster of four structures. Structural models show the conservation of the core aquaporin fold with the predicted backbone conformations of the models superimposing well with the recently solved 1.8 Å crystal structure of the rice Lsi1 NIP III silicic acid channel (32), with the highest similarity observed with the NIP4;1 and NIP6;1 models (RMSD =1.5 to 1.7 Å Table S2). These were analyzed further as examples of NIP I and II protein pores, respectively.

Examination of the NIP6;1 model predicts a more hydrophobic ar/R selectivity region compared to Lsi1 (Fig 1C). Unlike the invariant serine residue found at the H5 position in NIP III channels, NIP6;1 possesses a branched isoleucine residue at this position. Additionally, the invariant glycine residues found at H2 and LE_1_ in NIP III are occupied by alanines in NIP6;1. Despite these differences, the NIP6;1 pore, similar to Lsi1, is still wider than classical aquaporins (Fig. 1C) and allows access of a fifth residue (glycine 96 from helix 1 in NIP6;1) to the selectivity filter. In addition, residues involved in the binding and positioning of bound waters (Cys 39 and Ala 110 in NIP6;1, Fig. S2A) that are involved in metalloid transport in Lsi1 are conserved in the NIP 6;1 model. Thus, the predicted NIP6;1 pore model could potentially accommodate the additional bound waters at the selectivity filter that are a hallmark of the NIP III structure (Fig. 1C).

Based on the positions of the selectivity filter-associated water molecules in the Lsi1 structure, Saitoh et al. (32) modeled a silicic acid hydroxide interaction site (Fig. 1B). By using a similar approach with the waters within the NIP6;1 model, a potential interaction site for boric acid was investigated. Compared to the tetrahedral silicic acid molecule, boric acid is a smaller molecule with three hydroxyl groups oriented in a trigonal and planar arrangement. A model of boric acid docked into the NIP6;1 selectivity filter is shown in Fig. 2. Similar to the Lsi1 Si-bound model, potential hydrogen bond interactions with some selectivity filter residues, notably the conserved arginine and H1 (Gly96) and LE_1_ (Ala247) positions, are retained in NIP6;1. However, the replacement of the conserved Lsi1 Ser with Ile 238 in NIP6;1 at the H5 position would cause a steric restriction of the pore as well as a loss of hydrogen bond donor capability, consistent with the critical role of serine in providing silicic acid specificity (32,35).

**Figure 2.**
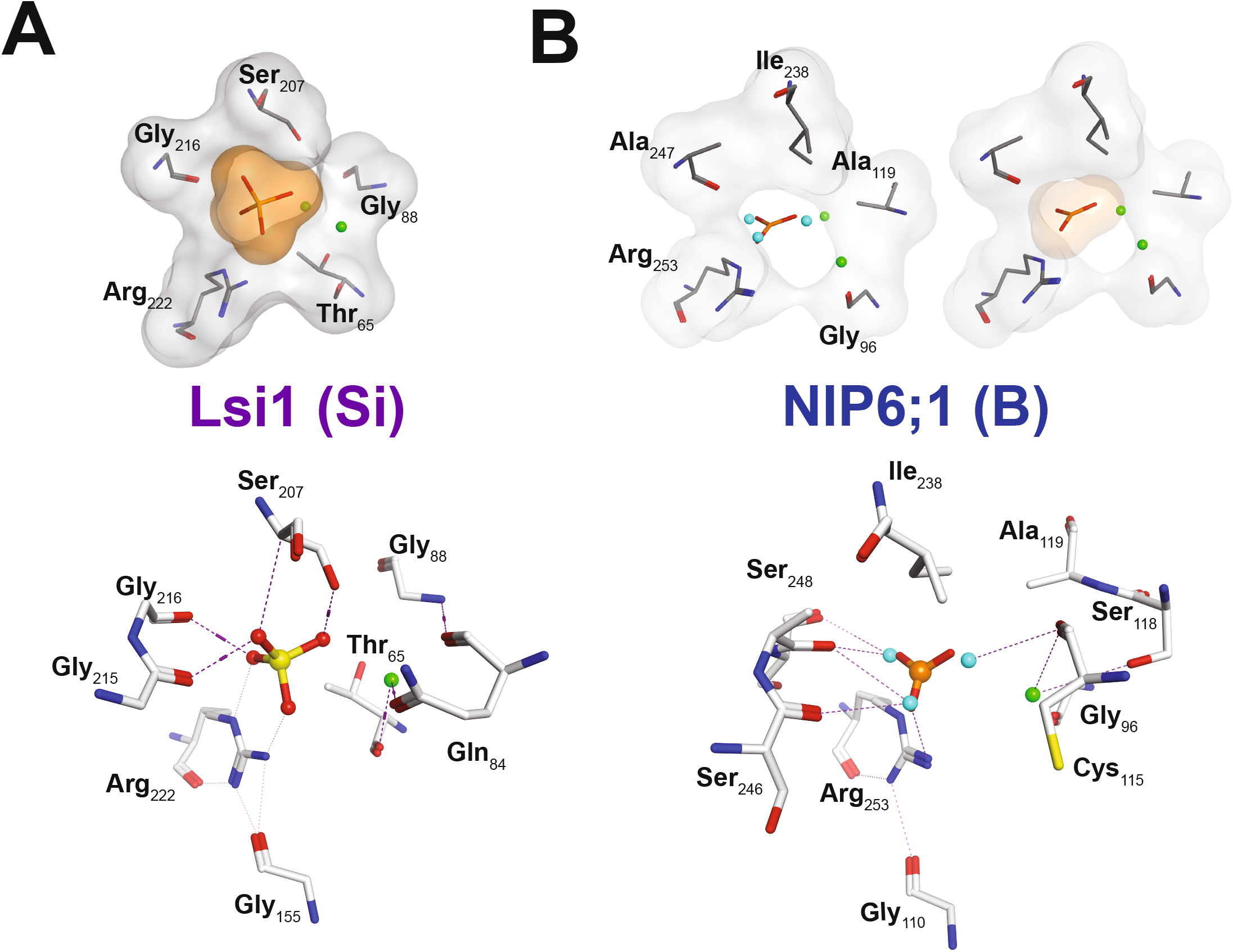
Boric acid bound model of NIP6;1. **A.** *Top*, the Lsi1 structure with Si(OH)_4_ bound at the selectivity filter taken from the QM/MM model of Saitoh et al. (32) with molecular surfaces of substrate and ar/R residues shown. *Bottom*, hydrogen bond contacts between Lsi1 amino acid residues and silicic acid (32). *Right*,. **B.** *Top Right*, model for NIP6;1 with boric acid docked to the ar/R selectivity filter. *Left*, The position of the five ar/R residues and the immobile (green) and selectivity filter bound (blue) waters from Fig. 1C are shown relative to the docked boric acid. *Bottom*, the position of predicted hydrogen bond contacts between NIP6;1 and boric acid. The position of immobile water 3 is shown in green, and three selectivity filter water molecules relative to the position of the boric acid hydroxyls are shown in blue.

The NIP4;1 model illustrates that NIP I protein pores are even more constricted and hydrophobic than the other NIP pore subtypes (Fig. 1C; Fig. S2). Like all NIP I pores (9,14), NIP4;1 possesses a conserved tryptophan at the H2 position which contrasts with the smaller glycine and alanine residues characteristic of NIP III and NIP II ar/R regions. To accommodate this bulkier residue, the H1 position is an invariant glycine in NIP I pores. Nevertheless, the result is a smaller ar/R with four residues instead of five and a calculated loss in the ability to accommodate the bound waters (water 3 and 9) and one of the transported waters (water 2) found in the Lsi1 channel (Fig. 1C). As a result, the remaining selectivity filter waters in NIP 4;1 are calculated to adopt a single file arrangement (Fig 1C). In this regard, NIP4;1 resembles microbial and animal aquaglyceroporins that have similar amphipathic ar/R selectivity filter properties with a narrow ar/R constriction that permits single file transport of water and solutes.

### 2.2 NIP I and II and corresponding H2 mutants show distinct metalloid uptake in oocytes

One result from the modeling is that the nature of the H2 residue determines the size and characteristics of the selectivity filter of NIP I and II proteins. To determine experimentally whether this residue affects the permeability metalloid hydroxide nutrients (B) and toxins (As), the boric acid and arsenous acid permeabilities of representative NIP I (NIP4;1) and II (NIP6;1 and NIP5;1) proteins, and their corresponding trp (NIP I-like) or ala (NIP II-like) H2 mutants, were evaluated in *Xenopus* oocytes (Fig. 3). *Xenopus* oocytes that express Arabidopsis NIP II and NIP I proteins show 20-fold greater As(OH)_3_ uptake rates compared to negative control oocytes, with no statistical differences in the permeability properties of NIPs from either pore subclass (Fig. 3A). Further, substitution of an alanine for tryptophan at the H2 position in the NIP I proteins (NIP4;1_W82A_), which would be predicted to increase the aperture of the ar/R, results in no significant differences in As(OH)_3_ uptake compared to wild type controls. Conversely, the substitution of a tryptophan for alanine in the NIP6;1 ar/R, which would be predicted to restrict the ar/R diameter, also showed no effect on As(OH)_3_ uptake rate (Fig. 3A). However, the NIP5;1_A117W_ mutant is an outlier that showed As(OH)_3_ uptake rates that were indistinguishable from negative control oocytes, reflecting the loss of transport function. Subsequent analysis (Fig. S4) showed that NIP5;1_A117W_ is impermeable to all substrates tested, and genetically is unable to complement *nip5;1-1* boron sensitive mutant in Arabidopsis screens. Therefore, for reasons that are not apparent, the NIP5;1_A117W_ mutation produces an inactive channel, and further comparisons focused on NIP4;1 and NIP6;1 and their mutants.

**Figure 3.**
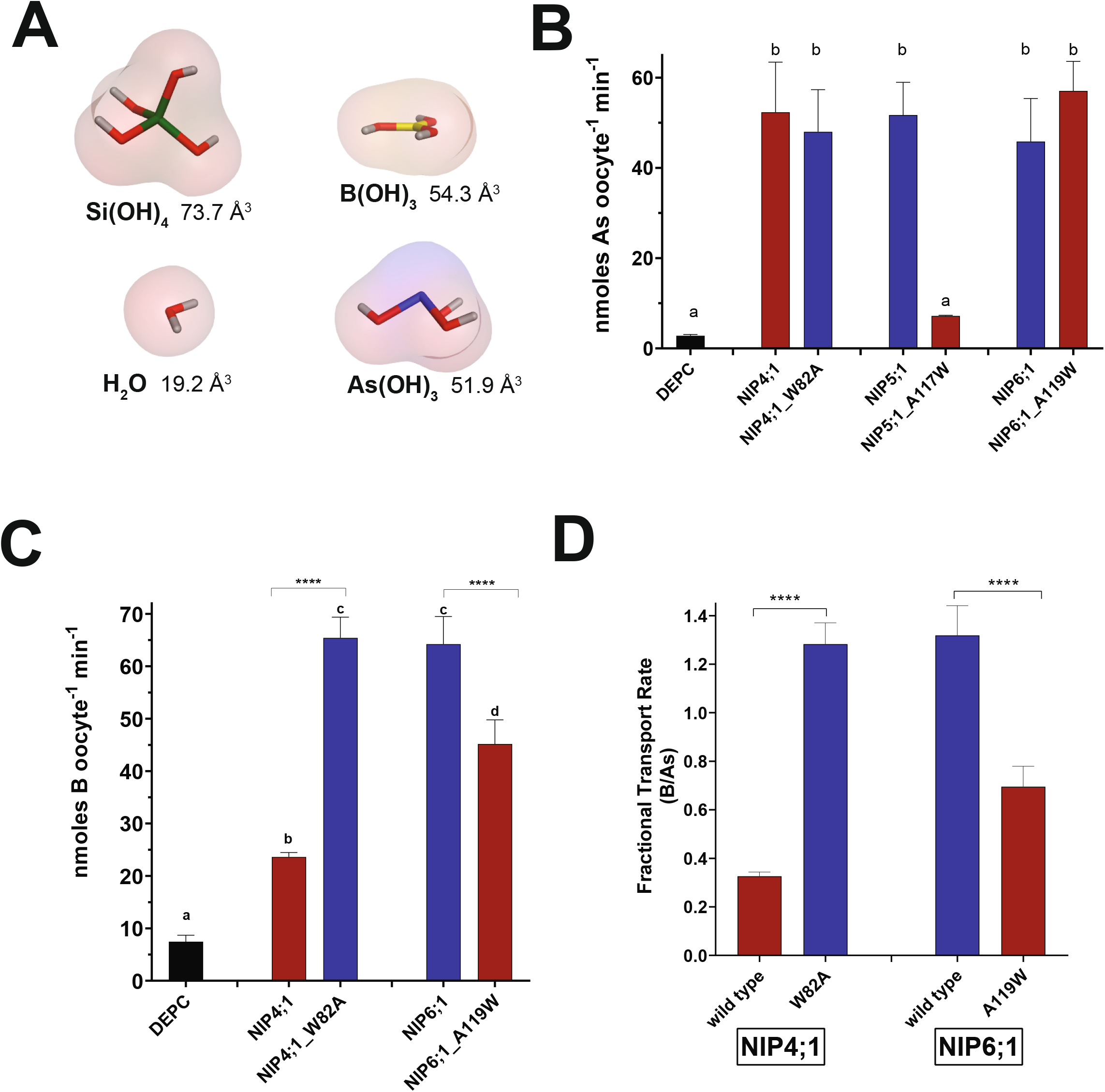
Comparison of the metalloid hydroxide permeabilities of NIP I and II wild type proteins and H2 mutants. **A.** Structural models of various metalloid hydroxides that permeate NIP channels illustrating their disparate geometries and sizes. Models were constructed and minimized in MOE and the molecular volumes were calculated by using VEGA ZZ (84). The C3 conformation of arsenous acid based on Ramírez-Solís et al. (43) is shown. **B.** Oocytes expressing GFP fusions of the indicated NIP or NIP mutants were subjected to As(OH)_3_ uptake analyses and As content was determined by ICP-MS. Values represent means ± SD (n = 4 pools of eight oocytes per pool, error bars indicate SD). DEPC represents negative control oocytes. **C.** Oocytes injected with the indicated NIP construct were subjected to boric acid uptake analyses and B content was determined by ICP-MS. **D.** Standardization of B uptake rates as a fraction of As uptake under the same conditions. The background (DEPC) oocyte uptake rate of B or As was subtracted, and the NIP-mediated B uptake is expressed as a fraction of the As uptake (n = 7 pools of 8 oocytes per pool, error bars show SD). Statistical significance in B, C, and D was determined by One-way ANOVA with a different letter indicating statistical significance (p<0.05). In panel C and D, ****, p<0.0001.

Next, the boric acid permeability of NIP4;1 and NIP6;1 proteins and their H2 mutants were compared. Since NIPs show indistinguishable arsenous acid permeability, for comparative purposes the boric acid permeability was normalized to the arsenous acid uptake. This allows the elimination of nonspecific background uptake of the two substrates as well as elimination of any differences related to slight variations in protein expression. Wild type NIP6;1 shows robust transport of both metalloids with a preference for B over As (fractional B/As permeability of 1.32) (Fig. 3D). In contrast, NIP4;1 was a poorer boric acid transporter with a 4-fold lower permeability (B/As = 0.32) (Fig. 3D). The NIP4;1_W82A_ mutant has substantially higher boric acid permeability (Fig. 3C), with a B/As permeability (1.28) statistically indistinguishable from the NIP6;1 boric acid channel. Conversely, the substitution of the H2 alanine in NIP6;1 with a tryptophan residue (NIP6;1_A119W_) results in a selective reduction in boric acid permeability (B/As = 0.69, Fig. 3D). Overall, the transport data show that the presence of the conserved NIP II-like alanine residue at the H2 position enhances the selective boric acid permeability in the NIP6;1 and mutant NIP4;1_W82A_ channels while exhibiting no influence on the permeability of a separate metalloid, arsenous acid.

### 2.3 NIP I proteins partially rescue the B deficiency phenotype of nip5;1 mutant seedlings

To evaluate further the ability of NIPI proteins as transporters of boric acid in plants, their ability to complement the low B sensitivity phenotype of the *nip5;1-1* T-DNA lines (36) was assessed. Transgenic plants that express the C-terminally tagged GFP fusions of various Arabidopsis NIPs and their H2 mutants under the control of the 35S promoter were generated in the *nip5;1-1* knockout background (*35S_pro_:NIP* transgenic plants). Western blot analysis of the seedlings of T2 transgenic lines showed the robust expression of the GFP fusion proteins (Fig. S3B), and confocal microscopy showed co-localization of the GFP signal with the fluorescent plasma membrane marker FM4-64 (Fig. S3C).

To determine whether the NIP I transgenes complemented the *nip5;1* phenotype, growth under sufficient (50 μM) and limiting boric acid (1 μM) was compared. Under sufficient boric acid conditions all plants, regardless of genotype showed normal growth (Fig. S5 and Fig. 4A) and were indistinguishable from wild type Col-0 plants. This is consistent with previous observations of *nip5;1-1* plants (36), and further indicates that overexpression of the NIP transgenes does not affect plant growth under these conditions. Under limiting B conditions, *nip5;1-1* plants showed defective growth that was complemented to different degrees by the various NIP I protein transgenes (Fig. S6, Fig. 4). Seven-day old seedlings from complementation lines expressing wild type *NIP1:1-GFP, NIP1;1_W94A_* and wild type *NIP4;1-GFP* constructs showed significantly longer root lengths and higher seedling fresh weights compared to *nip5;1-1* control plants. However, these constructs did not complement growth defects as well as the *NIP5;1* complementation line and showed significantly reduced root length and seedling weights compared to Col-0 wild type controls (Fig. S6). At later growth stages complementation lines with *NIP1;1* and *NIP4;1* constructs showed restored rosette leaf growth compared to *nip5;1-1* controls, but showed a delay in bolting compared to Col-0 plants and *NIP5;1* complementation plants (Fig. 4). B uptake analysis of all type 1 NIP complementation lines showed significantly enhanced B uptake and incorporation into rosette leaves compared to *nip5;1-1* negative controls, but B accumulation was significantly below Col-0 positive controls (Fig. 4C). Overall, the data show that NIP I proteins are able to facilitate boric acid uptake at a level that allows partial complementation of the *nip5;1* phenotype.

**Figure 4.**
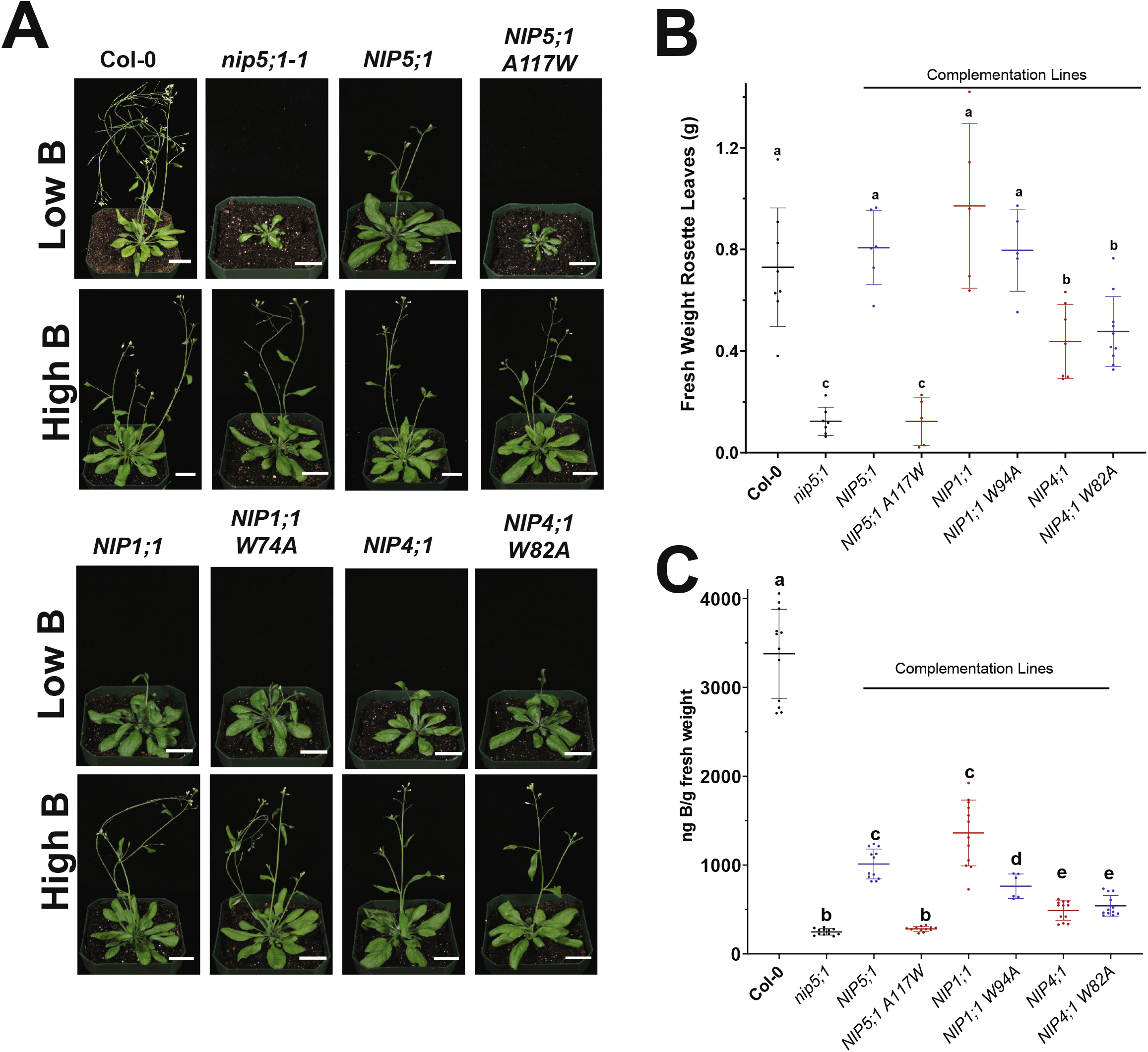
Complementation of *nip5;1* growth and B uptake phenotypes by NIP1 wild type and H2 ar/R mutant constructs. **A.** Representative plants (Col-0, *nip5;1-1* and *nip5;1-1* complemented with the indicated *NIP-GFP* fusion transgene grown for 40 days under limiting (**low B**) or sufficient (**high B**) boric acid conditions. Scale bar = 2cm. **B.** Fresh weights of rosette leaves (each data point represents a separate determination) collected from 40 day old plants grown under low B conditions as in Panel A. **C.** Plants were grown under low B conditions, and B uptake analysis was done as described in the Materials and Methods. For panels B and C, statistical analysis was performed by One-way ANOVA with different letters indicating statistical significance (p<0.05).

### 2.4 The watertight property of NIP II proteins

The NIP III Lsi1 structure reveals that the pore is unusually hydrophilic and hydrated compared to other members of the aquaporin family (32), with 16 water molecules occupying each monomeric channel (Fig. S1B), and in general NIP III proteins possess strong aquaporin activity in addition to its ability to transport silicic acid and other metalloids (10). The NIP6;1 model also shows a wide selectivity filter that is able to accommodate the three transported waters within the ar/R region as well as two bound waters found in the Lsi1 structure (Fig. 1C), While the structural model predicts that NIP II proteins should be able to accommodate water movement, examination of NIP6;1 and NIP5;1 in *Xenopus* oocytes show that they lack detectable aquaporin activity and are statistically indistinguishable from negative control oocytes with respect to water permeability (Fig. 5). In contrast, NIP I proteins, which have a more restricted selectivity filter (Fig. 1C), show substantial aquaporin activity (Fig. 5). This difference between NIP I and NIP II aquaporin activities is controlled by the nature of the H2 residue in the selectivity filter. Substitution of the canonical H2 tryptophan in NIP I proteins with the NIP II-like alanine residue (NIP1;1_W94A_, NIP4;1_W82A_, nodulin 26_W77A_) significantly diminishes or abolishes water permeability (Fig. 5). Conversely, substitution of the tryptophan for alanine in NIP6;1 results in strong gain of function aquaporin activity (Fig. 5).

**Figure 5.**
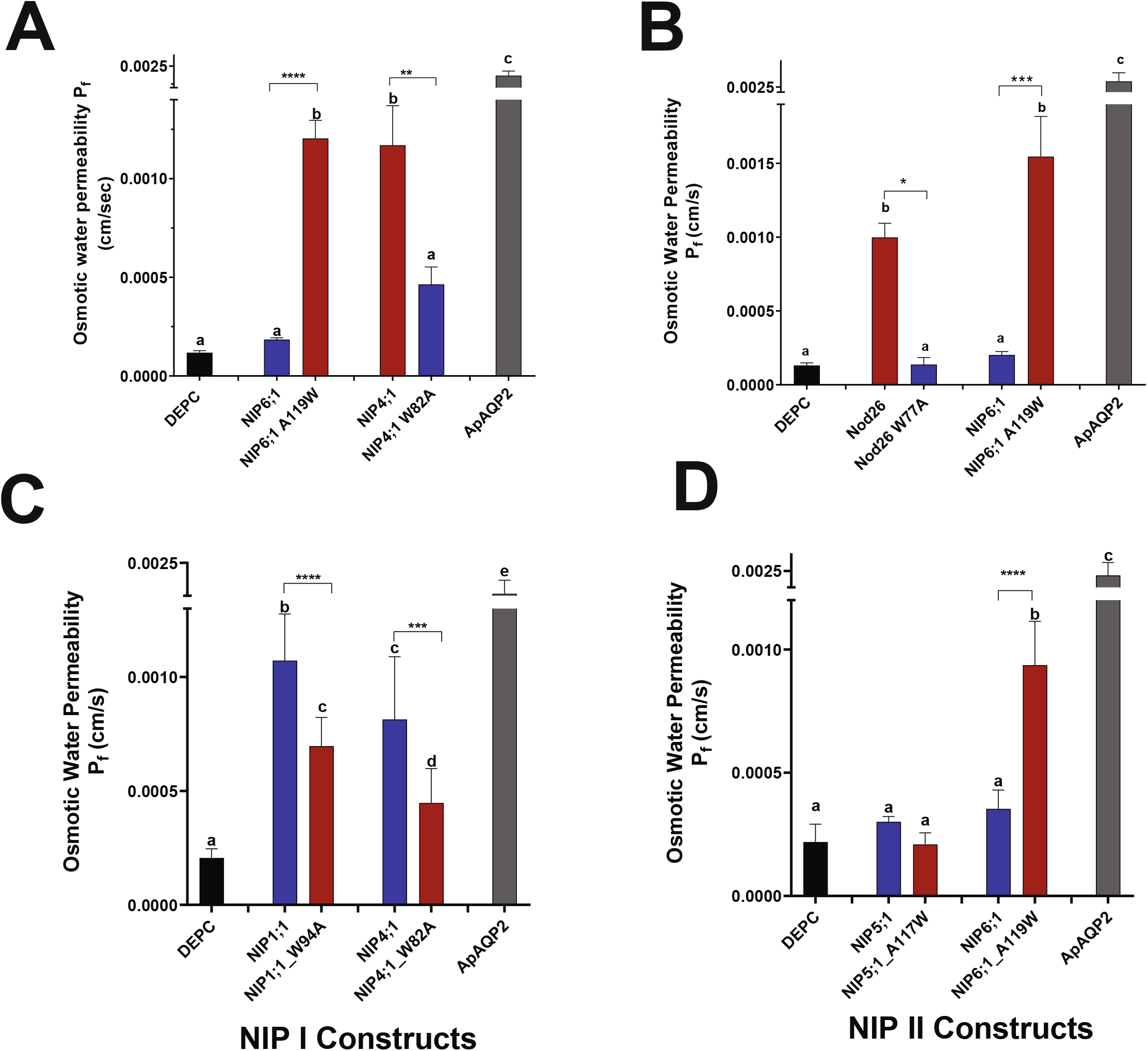
Comparison of the aquaporin activities of NIP I and II wild type proteins and H2 mutants. **A.** Comparison of the osmotic water permeability of oocytes injected with cRNA encoding flag-tagged wild type Arabidopsis NIP6;1 and NIP4;1, or corresponding H2 ar/R mutants (NIP6;1 A119W and NIP4;1 W82A). The osmotic water permeability (P_f_) was determined from the rate of oocyte swelling upon incubation in a hypoosmotic Ringer’s solution (46). Values represent means ± SD (n = 8-24 oocytes for each sample). DEPC, negative control oocytes injected with sterile DEPC water. ApAQP2, positive control oocytes injected with the flag-tagged *Acyrthosiphon pisum* AQP2 aquaporin (62). **B.** Comparison of osmotic water permeability of NIP I proteins (NIP1;1 and NIP4;1) and their corresponding tryptophan to alanine H2 mutants. Oocytes (n=16-20) were injected with cRNA encoding the indicated protein with an in-frame C-terminal GFP tag. **C.** Comparison of osmotic water permeability of NIP II proteins (NIP5;1 and NIP6;1) and their corresponding tryptophan to alanine H2 mutants. Oocytes (n=8-10) were injected with cRNA encoding the indicated protein with an in-frame C-terminal GFP tag. Statistical significance for each data set was determined by One-way ANOVA with a different letter indicating statistical significance (p<0.05). Asterisks indicate statistical significance between wild type and H2 mutant pairs for each NIP subtype determined by pairwise t-test (**, p<0.01; ***, p<0.001; ****, p<0.0001).

Why are NIP II proteins unable to transport water like their NIP I and NIP III counterparts? One possible explanation comes from the examination of the different clusters of the NIP6;1 structural models generated from Alphafold2/MD simulations that show distinct orientations (“up” vs “down”) of the canonical ar/R arginine (Arg253) relative to the pore axis (Fig. S8). In the “up” conformational model, the arginine is oriented towards the extracellular side of the membrane, parallel to the pore axis, and is stabilized by a hydrogen bond with the backbone carbonyl of Gly186 in the central C-loop (Fig. S7). This is similar to the orientation of the canonical ar/R arginine in most aquaporin structures and represents an open configuration. In the “down” configuration, Arg253 hydrogen bonds with Thr97, a conserved residue of NIP II proteins found in transmembrane helix 1 (Fig. S7; Fig. 6A). This configuration places the arginine side chain within the pore axis in a position that could affect water transport. Examination of the structural clusters of NIP I models and NIP6;1_A119W_ mutants show that the selectivity filter arginine remains in the “up” open configuration throughout the MD simulation trajectory (Fig. 6A), possibly due to the steric constraints of the bulkier tryptophan side chain.

**Figure 6.**
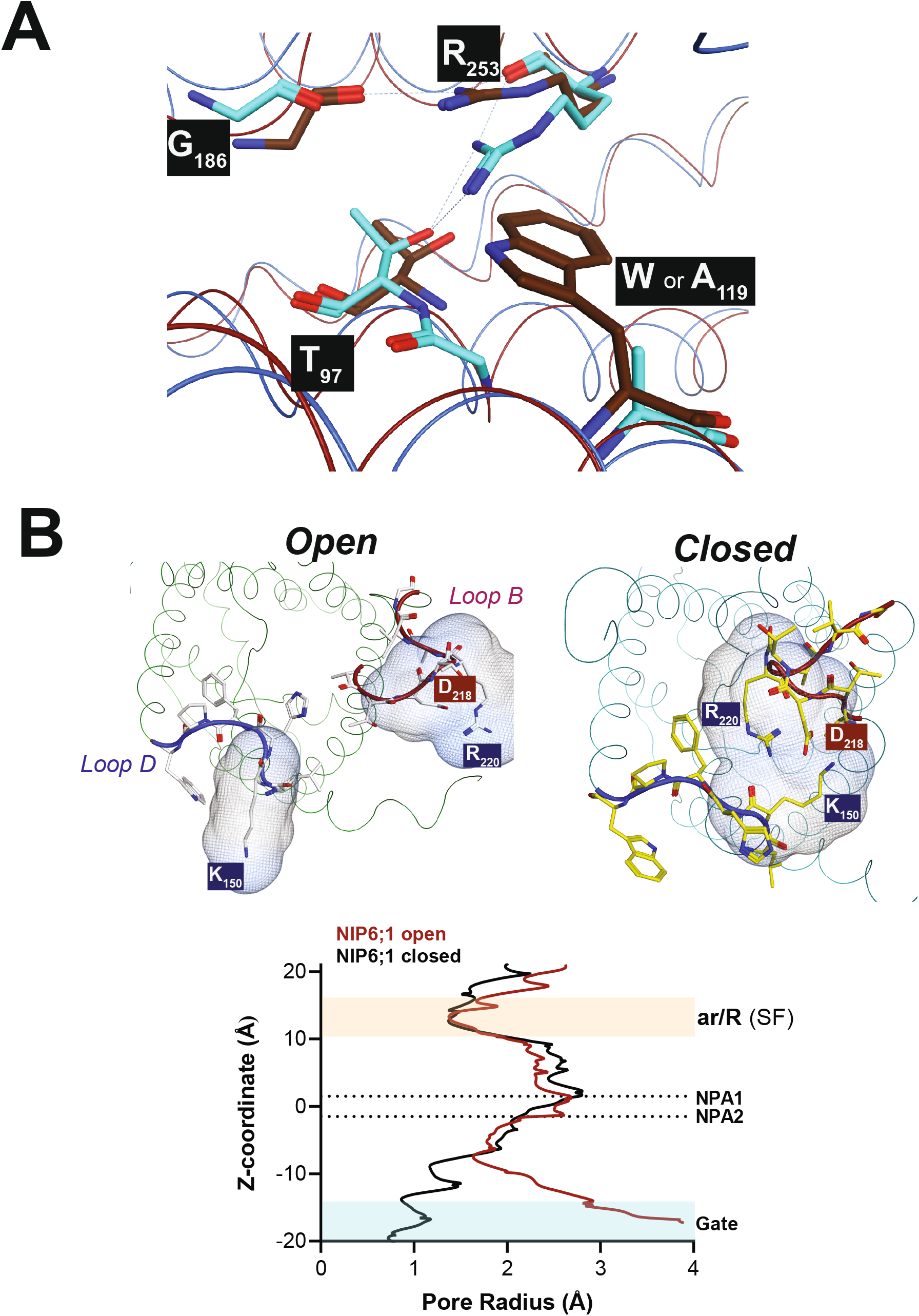
Potential gating residues in the NIP6;1 model. **A.** Comparison of the selectivity filter arginine orientation in the NIP6;1 “down” conformation and the NIP6;1A119W mutant homology model. The down configuration is not observed during MD simulations of the the mutant model, presumably due to the steric constraints of the bulkier tryptophan sidechain. **B. *Top***, Models for the open and closed states for NIP6;1. ***Bottom*** Plot of the calculated pore diameter along the Z-pore coordinate of the two NIP6;1 models. The data were obtained by using Mole*Online* 2.5 (83) in the Channels mode. The position of the conserved NPA motifs that define the center of the pore as well as the ar/R selectivity filter region are indicated.

Another consideration is the potential gating of solute and water access to the pore by interhelical loop regions based on a comparison of the two recently solved structures of Lsi1 in an open conformation (pore filled with waters, Saitoh et al. (32)) and a closed structure (van den Berg et al. (33)). The closed structure results from a reorientation of loop D so that it forms electrostatic/hydrogen bond interactions with loop B, essentially blocking the pore access at the intracellular vestibule (Fig. S8B). Examination of the sequences of NIP II proteins (Fig. S1C) shows conservation of the potential gating residues in the loop D and loop B region. A comparison of NIP6;1 structural models shows that protein could occupy an open or closed state stabilized by similar interactions observed in the closed Lsi1 structure (Fig. 6B). Such a structure would effectively restrict the pore diameter to less than that of a water molecule (Fig. 6C) which would prevent water movement.

## 3. Discussion

### 3.1 Distinct metalloid permeability of NIP channel subtypes

All members of the NIP family facilitate the transport of metalloid hydroxides (25). These include nutrients such as boric acid and orthosilicic acid, as well as toxic hydroxides of As (III), Ge and Sb. These compounds form weak Lewis acid structures with different hydroxyl group geometries ranging from trigonal planar (B[OH]_3_), to trigonal pyramidal (As[OH]_3_ and Sb[OH]_3_) to tetrahedral (Si[OH]_4_ and Ge[OH]_4_) molecules. While metalloid transport is a common feature of NIP proteins, the three NIP pore subtypes show distinct selectivity for various metalloids. The most selective are the NIP III pores which are specific for the tetrahydroxylated Si(OH)_4_ and Ge(OH)_4_ (15,35). From the elucidation of the Lsi1 structure, NIP III proteins contain an exceptionally wide and hydrophilic channel with a conserved five amino acid ar/R selectivity filter and two tightly bound waters that are hallmarks of the NIP III pore (32). These features accommodate the bulky Si(OH)_4_ molecule both sterically as well as enthalpically by providing several hydrogen bond contacts as the molecule traverses the pore (32).

Analysis of the NIP I and II pore models suggests two features that might hinder Si(OH)_4_ permeation. First, NIP I and II pores have a hydrophobic substitution (valine or isoleucine) for serine at the H5 position as well as a glycine for threonine at the H1 position (Fig.1C). This would reduce the hydrophilicity and hydrogen bonding capability of the selectivity filter. Second, the conserved glycines at H2 and LE_1_ in NIP III are replaced with larger sidechains in NIP I and NIP II which would restrict the diameter (particularly for NIP I) of the pore. As a result the larger Si(OH)_4_ (74 Å^3^) would be sterically restricted and would be unable to form optimal hydrogen bond contacts compared to the smaller As(OH)_3_ or B(OH)_3_ (52 to 54 Å^3^ respectively) molecules.

In contrast to Si(OH)_4_, As(OH)_3_ is a promiscuous substrate that is transported by all three functional NIP sub-classes (22–24,35,38–42). This is consistent with our observations that all NIP proteins tested here are equally permeable to As(OH)_3_ and that substitutions at the H2 position (tryptophan or alanine) have no effect on transport of this substrate. Unlike As(OH)_3_, comparison of Arabidopsis NIP6;1 and NIP 4;1 show that boric acid is more readily transported by the NIP II pore. To examine how the NIP6;1 selectivity filter could lead to this difference, a NIP6;1 model with boric acid docked into the selectivity filter was generated by using the hydrated structure of Lsi1 as a template (Fig. 2). Within the NIP6;1/boric acid structural model, the three boric acid hydroxyl groups are positioned within a single plane and form hydrogen bonds with the conserved ar/R arginine and backbone carbonyls or side chains from the H1 (glycine 96), LE_1_ (Ala 247) residues and adjacent residue Ser 248. Similar to the Lsi1 structure, the small residue at H2 (Ala 119) of NIP6;1 allows access for the H1 residue resulting in wider five amino acid ar/R. In the Lsi1 pore, such an arrangement leads to occupancy of the pore with two bound waters that participate in silicic transport in MD simulations (32). We hypothesize that a similar spatial arrangement in NIP6;1 may be required for optimal interaction with the trigonal boric acid hydroxyl groups (Fig. 2B).

In contrast, the NIP4;1 structural model contains a bulky tryptophan side chain in the H2 position that sterically prevents the access of the H1 residue to the ar/R region. This results in a classical arrangement of four amphipathic amino acids in the ar/R selectivity filter similar to bacterial and animal aquaglyceroporins. Compared to the NIP6;1 model, this would prevent the positioning of water 3, as well as sterically restrict the pore at one of the proposed positions of the boric acid molecule (occupied by water 2 in Lsi1 and NIP6;1 models). This prediction fits with the observation that boric acid is more poorly permeated in NIP4;1, and that the permeability of the protein for this metalloid increases four-fold with a gain-of-function substitution of alanine for tryptophan at the H2 position. Conversely, the substitution of a tryptophan for alanine in NIP6;1 has an opposing effect, causing a two-fold reduction in permeability.

The NIP I and II pore models do not explain why As(OH)_3_, which has a molecular volume that is similar to boric acid, permeates NIP4;1 and NIP6;1 equally. However, a closer comparison of the two metalloid hydroxides suggests that this may be due to differences in geometry, electronic properties, and amphiphilicity. Boric acid (Fig. 3A) is a planar trigonal molecule (120° O-B-O bond angle) whereas As(OH)_3_ is a trigonal pyramid with more restricted bond angles (O-As-O 97° based on X-ray absorption spectroscopy and Density Functional Theory calculations, 43). As a result, unlike the flat trigonal arrangement of the boric acid hydroxyls, the three hydroxyl groups in As(OH)_3_ are clustered to one side of the molecule, while the lone pair of electrons occupies the opposing side of the As atom. Based on quantum calculations and thermodynamic comparisons, this arrangement predicts that As(OH)_3_ has an amphipathic geometry (with high hydrophobicity on the lone electron pair side) that is not shared by boric acid (44), but is similar in properties to another common aquaporin substrate, glycerol (45). Notably, similar to As(OH)_3_, glycerol permeates NIP I and NIP II pores equally (46). Conversely, in addition to NIP proteins, other plant and animal aquaglyceroporins readily transport As(OH)_3_ (38,47). Overall, the findings confirm the notion that if an aquaporin fluxes glycerol, it will often also flux As(OH)_3_.

### 3.2 NIP II proteins as specialized, water-impermeable boric acid channels

NIP II proteins in dicots are phylogenetically segregated into three subclades with preferential expression in roots (NIP5;1), shoots (NIP6;1), and the anthers of developing flowers (NIP7;1) (48). Genetic and biochemical evidence shows that all three NIP II proteins : 1. Are plasma membrane-localized proteins that flux boric acid (36,48–50); 2. are tightly regulated and specialized at multiple levels (transcription, translation, polar localization, and trafficking) for boric acid uptake and transport under conditions of boric acid limitation to support the formation of the rhamnogalacturonan II (RG-II) pectin cell wall in collaboration with the BOR transporters (reviewed in 19–21); and 3. Are largely impermeable to water (14,26,36,42,46,49,50).

While the NIP II subclass is specialized for boric acid uptake and distribution, the biological functions of NIP I proteins as boric acid channels in Arabidopsis is less clear. While NIP I proteins such as NIP4;1 have reduced boric acid permeability compared to NIP II proteins, over expression does facilitate boric acid uptake and partial complementation of the boron-deficient phenotype of *nip5;1* plants. Diehn et al. (26) found similar results with the NIP I orthologs of *Brassica napus*, and suggested a potential tissue specific B-transport role for some this NIP class that could help coordinate B homeostasis in coordination with NIP II and BOR transporters. However, despite the observation that these proteins are permeable to boric acid to some level, additional genetic and physiological data is required to firmly support this as their primary biological function. In contrast, evidence for lactic acid transport (31), As uptake from soils (23,24), and aquaporin and ammonia-porin activities in nitrogen fixing symbioses (27,29,51) suggest additional biological functions for this NIP subclass.

The water-tight feature of NIP II proteins is intriguing, and one could ask from a biological perspective why aquaporin activity would be restricted in NIP II proteins? The answer may come from the predominant plasma membrane localization of these proteins. Unlike animal cells, the water permeability of the plant plasma membrane is tightly and coordinately regulated with other internal organellar membranes, particularly the tonoplast of the vacuole, to enable cell volume regulation and cytoplasmic buffering. For example, the water major aquaporin proteins of the plant plasma membrane (PIPs) are stringently regulated by gene expression modulation, reversible trafficking between internal membranes and the cell surface, as well as gating at the protein level through hetero-oligomerization, phosphorylation, pH and divalent metal cations (52–56). In the case of NIP II proteins, the ability to readily transport boric acid in a water-tight manner may be necessary to prevent disruption of cell turgor and cell volume regulation while enabling transcellular movement of this solute to tissues of need.

The lack of water permeability of NIP II pores is the result of a small amino acid, usually alanine, at the H2 position of the ar/R selectivity filter. This is counter-intuitive since modeling predicts that NIP II pores are wide enough to accommodate multiple water molecules. Conversely, the substitution of a large aromatic residue (tryptophan), which restricts pore diameter, opens the pore to water transport. Based on our modeling results, we hypothesize two potential mechanisms (“pore pinching” or “loop capping” by the terminology of Hedfalk et al. (57)) that could regulate NIP6;1 pore access and water or solute permeability. With respect to the first mechanism, MD simulation of our NIP6;1 models shows that the ar/R arginine occupies two rotameric states that could restrict access of transported substrate to the selectivity filter. The substitution of tryptophan predicts the orientation of arginine preferentially in an “up” configuration that would open the pore. A similar case exists for the *E. coli* aquaporin AQP Z, with the crystal structure of the protein revealing arginine rotameric states, stabilized by similar hydrogen interactions as in NIP6;1, that open or close the pore to water movement (58,59).

The loop capping model is postulated based on alternative open and closed conformations observed in crystal structures of Lsi1 (32,33). The closed conformation is stabilized by electrostatic and hydrogen bond interactions between acidic and basic amino acid side chains in loop D and loop B on the cytosolic side of the membrane (33). During MD simulation, reorientation of loop D is observed and both open and closed conformations were occupied. Calculation of bidirectional water flux of the two conformers showed that the closed configuration has substantially reduced water permeation, suggesting that Loop D could potentially serve as a gate for water and solute transport (33). Interestingly, the features of loop D that stabilize the closed conformation are conserved across all NIP protein subclasses suggesting this is a common feature of the subfamily (cf. Fig. S1 and Fig. S8B). Similar regulation of gating in loop D of other members of the aquaporin family has been documented (60). In plants, a notable example is provided by the SoPIP2;1 protein in which loop D forms a cap over the pore aperture (61). Regulation and channel opening occur through several mechanisms including pH, phosphorylation and calcium ion interaction (61). The aquaporin activity of some NIP proteins (e.g., soybean nodulin 26), are regulated by phosphorylation (51), and the potential regulation of the pore properties of NIP proteins by these mechanisms merits further structural evaluation.

## 4. Experimental Procedures

### 4.1 Molecular Cloning

cDNA constructs with NIP coding sequences (CDS) were generated from total RNA isolated from *Arabidopsis thaliana* Col-0 seedlings (*NIP1;1*, *NIP5;1* and *NIP6;1*) or flowers (*NIP4;1*) as previously described (46,49,50). The CDS encoding the aquaporin control protein, *Acyrthosiphon pisum* AQP2, was generated as described in Wallace et al. (62). GFP fusions of the NIP and AQP constructs were generated by the method described in Beamer et al. (31). Briefly, the NIP or AQP CDS were amplified by using gene specific primers (Table SI) and were cloned into the BioVector Gateway entry vector Fu28 (ABRC stock no: CD3-1822; Wang et al., 2013), which contains an eGFP CDS at the 3’ end of a multiple cloning site. The primers were designed with a linker of three glycines between the NIP and eGFP coding sequences. A gateway LR reaction (Invitrogen) was performed to transfer the CDS-eGFP construct from the entry clone to the Gateway destination vectors for *Xenopus laevis* expression (pT7TSGW) or for plant transformation for complementation experiments (pB7WG2; Karimi et al. (64)). For *Xenopus* experiments, the destination vector pT7TSGW was generated by incorporating the Gateway cassette system from pB7WG (from VIB-UGent Center for Plant Systems Biology, Ghent University, Belgium) into the *Xenopus laevis* expression plasmid pT7TS (65) at the *EcoR*V site.

Site-directed mutagenesis to generate alanine or tryptophan substitutions at H2 positions in NIP constructs was done by using the mutagenesis primers listed in Table SI and the Q5 Site-Directed Mutagenesis Kit (NEB) as described previously (50). Flag-tagged *Xenopus* expression constructs were generated in the pXβG-ev-1 vector (46). The sequence of all constructs were confirmed by Sanger sequencing with a Perkin Elmer Applied Biosystems 373 sequencer at the University of Tennessee Molecular Biology Research Facility (Knoxville, TN, USA).

### 4.2 Seedling growth, B phenotype analysis, and B uptake in plants

All *in planta* experiments were conducted in the Col-0 ecotype of *Arabidopsis thaliana*. The *nip5;1-1* T-DNA insertion line (Salk_122287C, Takano et al. (36)) was a kind gift of Professor Junpei Takano, Osaka Prefecture University. For seedling growth under limiting or sufficient boron conditions on plates, MGRL media (37) with 1% (w/v) sucrose was treated overnight with 3g/L of Amberlite IRA743 (Sigma) resin to remove residual boron (66). Boric acid was then added to a final concentration of 1 μM or 50 μM, in addition to 1.5% (w/v) Phytagel (Sigma), before sterilization. Seeds were sterilized in 8% (v/v) sodium hypochlorite, 0.05% (v/v) Tween-20 for 15 min and were washed five times with sterilized deionized water. Seeds were sown in the MGRL medium with 1μM B, were vernalized at 4°C for 2 d, and were grown vertically with a long day (LD) cycle of 16 hours light (100μmol m^−2^ s^−1^) and 8 hours dark. After 7 days, the plants were imaged for GFP signal using a wide-field epifluorescence microscope (DM6000 B; Leica). Images were collected with a DSLR camera (Canon Rebel XS) and seedling weight and root lengths were recorded.

For growth in soil, transgenic lines were sown onto MGRL media supplemented with 0.1% (w/v) sucrose, 30μg/mL BASTA and 50 μM B, and were vernalized and grown under LD conditions as described above. After 10 days the plants were transferred to soil, were returned to LD conditions, and were watered every 5 days with MGRL media supplemented with either 0.3 μM or 30 μM boric acid.

For the determination of boric acid uptake and content of plants, rosettes from 6-12 independent 40-day old plants were harvested, extracted and analyzed by the method of Diehn et al. (26). Samples were digested in nitric acid at 70°C overnight, and the B content was determined by ICP-MS using an Agilent 7500 cx instrument operating in the He collision mode (Spectroscopy and Biophysics Core, University of Nebraska, Lincoln, NE, USA). The samples were diluted 10 to 20-fold in 2% (v/v) HNO_3_ and were supplemented with 50ppb ^71^Ga as an internal standard.

### 4.3 Generation of Arabidopsis complementation lines

Transgenic complementation of *nip5;1-1* mutant lines was done with Arabidopsis *NIP-GFP* translational fusion constructs under the control of the 35S CaMV promoter by the general method described in Beamer et al. (31). *Agrobacterium tumefaciens* GV3101 strains carrying the desired *CaM35S_pro_:CDS-eGFP* construct was used to transform *nip5;1-1* background using the floral dip method (67). The transgenic lines were identified by initial screening and selection on MS media supplemented with 15 μg/mL BASTA. For confirmation and genotyping, genomic DNA was extracted from 2-wk old seedlings by using the *Wizard* Genomic DNA purification kit (Promega) and was subjected to PCR analysis by using a primer set consisting of a specific CaM35S promoter sequence and a gene-specific primer (Table S1). Seeds from homozygous T2 generation complementation lines were used for all analyses in this study.

### 4.4 Expression and functional analyses in Xenopus laevis oocytes

*Xenopus laevis* expression and functional analyses were performed as previously described (46,49,50). NIP or AQP expression constructs were linearized by digestion with *Xba*I, and capped cRNA was generated by in vitro transcription using the mMessage mMachine kit (Ambion). Stage V and VI *Xenopus* oocytes were collected surgically and defolliculated (51) or were obtained from Ecocyte Bioscience (Austin, TX). Oocytes were microinjected with 46nL of 1μg/uL of cRNAs or with an equivalent volume of RNase-free water (DEPC) as a negative control using a “Nanoject” automatic injector (Drummond Scientific Co., Broomall, PA, USA). Prior to assay, the oocytes were cultured for 72 h at 16°C in standard Ringer’s solution (96mM NaCl, 2mM KCl, 5mM MgCl_2_, 5mM HEPES-NaOH pH 7.6, 0.6mM CaCl_2_, 190 mosmol/kg) supplemented with 100μg/mL penicillin/streptomycin.

The boric permeability of the oocytes was determined as described previously (49) by incubation of groups of eight oocytes in standard Ringer’s solution supplemented with 2 mM boric acid. Assays were conducted at 16°C for 20 to 30 min, after which oocytes were washed five times with ice-cold (1mL per 8 oocytes per wash) standard Ringer’s solution without boric acid, followed by homogenization and overnight digestion at 65°C in 100μL of nitric acid. The uptake of arsenous acid was done identically except in the presence of sodium arsenite instead of boric acid. The As or B content of the digests were determined by ICP-MS analysis as described above.

The osmotic water permeability (*P_f_*) of the oocytes was measured by the standard swelling assay described previously (68). Oocytes were transferred from standard Ringers to hypoosmotic Ringer’s media (60 mosmol/kg) and the rate of oocyte swelling ((d*V/V_o_*)/dt) was determined from the cross-sectional area change determined by video microscopy. The *P_f_* was calculated from:

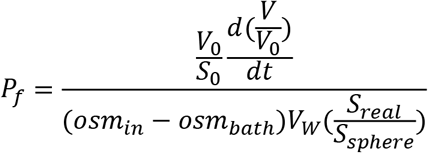

Where V is the volume at a specific time; V_o_ and S_0_ are the initial oocyte volume and cross sectional area; osm_in_ – osm_bath_ is the osmotic gradient; V_W_ is the partial molar volume of water, and S_real_/S_sphere_ is a surface area correction constant that accounts for the topology of the oolemma (69).

### 4.5 Other Analytical and Microscopy Techniques

For detection of NIP-GFP protein in oocytes, SDS-polyacrylamide gel electrophoresis (SDS-PAGE) and Western blot analyses were performed using 10μg protein from oocyte lysates. For the analysis of NIP-GFP expression in complementation lines, 10 d old seedlings were ground in liquid nitrogen and were extracted with SDS-PAGE sample buffer (70). Rabbit anti-GFP polyclonal antibodies (Abcam) were used as the primary antibody for the detection of NIP-GFP.

For GFP localization, 7 d old seedlings grown were stained with 4μM FM4-64 for 10 min prior to imaging with a Leica SP8 white laser system as described by Beamer et al. (31). The emission filter was set to 495 to 550 for GFP, 580 to 650 for FM4-64 and 680-720 for chlorophyll.

### 4.6 Computational methods and protein modeling techniques

The NIP structural models were obtained using Alphafold2, which is an AI system developed by DeepMind to predict the 3D structures of proteins from their amino acid sequences (34,71,72). Models of the mutants were generated using the Molecular Operating Environment (MOE, 2022). The NIP structural models were refined by molecular dynamics (MD) simulation performed using Amber simulation engine (73). To prepare the protein structures for MD they were hydrated using the water model TIP3P (74) in an octahedral box of at least 10 Å around the protein in each direction. The MD was performed using Amber14 ff14SB (75) force field with a non-bonded cutoff of 10 Å using the Particle Mesh Ewald algorithm (76). Restraints were applied following the procedure of Dutagaci et al. (77) for the structural refinement of membrane proteins.

All systems were initially energy minimized while applying restraints on water and ions with a 5 kcal/mol/Å^2^ force constant. The systems were energy minimized with 5000 steps of steepest descent followed by 5000 steps with the conjugate gradient method. The systems were further equilibrated using restraints on Cα and Cβ atoms with a force constant of 0.5 kcal/mol/Å^2^ followed by heating to 300K, and 1000 MD steps were performed. The SHAKE algorithm (78) was used to constrain all bonds involving hydrogen in the simulations. MD production runs were performed at 300K using the NPT ensemble and a 2 femtosecond time step. The temperature was fixed with the Langevin dynamics thermostat (79) and the pressure was fixed with the Monte Carlo barostat (80). Cα atoms were restrained during the simulations with a force constant of 0.025 kcal/mol/Å^2^ to avoid large deviations from the initial structures. Thirty nanosecond production runs were performed for each system. The structures obtained during the last 10 ns of each of the simulation trajectories were segregated into four clusters based on the root mean square deviation (RMSD) of the backbone atoms using the hierarchical agglomerate clustering algorithm present in the Cpptraj module (81). This method helps in dimension reduction and generates an ensemble of structures that also takes the local and global motion of the protein into account.

Comparison of the conformations and RMSD calculations of the structural models was performed with MOE. The dimension of the transmembrane pores along the z-coordinate of protein models was calculated by using the MOLEonline web server in the “channels” mode (Probe Radius 5, Interior Threshold 1.5, Merged Pores On) for the open structures and “Pore mode” (Probe Radius 5, Interior Threshold 0.3) for the closed structures (82,83). Models of metalloid hydroxides were constructed and energy minimized in MOE and the molecular volumes were calculated by using VEGA ZZ (84). The PDB coordinates for the open (7CJS, Saitoh et al. 32) and the closed (7NL4, van den Berg et al. 33) structures of the rice silicic acid channel Lsi1 were obtained from the NCBI Structure database, and the silicic acid bound Lsi1 structural model (Model 2) was taken from Saitoh et al. (32). Boric acid docked structures of the NIP6;1 protein model were constructed using MOE by using the selectivity filter waters of Lsi1 structure (7CJS, waters 1,2 and 4) as templates for orientation of the three hydroxyl groups of the boric acid molecule. In the final boric acid-bound NIP6;1 protein structure, the boric acid molecule was energy minimized to take into account the trigonal planar geometry of the hydroxyl groups. The energy minimization was performed in MOE by keeping the protein fixed with an RMS gradient of 0.1 kcal/mol/A^2^.

## Supporting Information

This manuscript contains supporting information.

## Data and materials availability

All data is available in the manuscript. Transgenic seed lines and DNA constructs will be made available from the corresponding author upon request.

## Acknowledgements

We thank Dr. Junpei Takano, Osaka Prefecture University for his generous gift of *nip5;1-*1 seeds. We are thankful to Drs. Andreas Nebenfuehr and Jaydeep Kolape (University of Tennessee, Knoxville) for assistance and advice on the confocal microscopic analyses.

## Author Contributions

**Z.B.**, organized and executed the majority of the experiments, and together with D.M.R. and P.R., wrote the manuscript; **P.R.**, directed the design and supervise the in planta experiments, assisted in generation of the *nip5;1* complementation lines, assisted in B and As uptake assays in *Xenopus* oocyte, involved in analyzing the data and organizing the manuscript; **R.A.**, generated the homology NIP homology models, carried out MD simulation experiments and substrate-docking analysis; **T.L.**, performed water transport assays in *Xenopus* oocyte; **K.G.**, undergraduate research student who assisted in *Arabidopsis* NIP-GFP localization as well as the *nip5;1* complementation experiments; **K.O.**, undergraduate research student who assisted in the screening of transgenic lines used in this study as well as early phenotype analysis for the *nip5;1* complementation experiments; **J.S.**, supervised the computational modeling and simulation experiments; **D.M.R**., supervisor of the project, obtained funding for the work, and was involved in the design of experiments, analysis and interpretation of the data, carried out the modeling of gated structures of NIP6;1 along with ZB and RA, and directed the writing and editing of the manuscript. DMR agrees to serve as the contact, communication, and source for materials emerging from this project.

