## Supplemental Figures for "Differential Boric Acid and Water Transport in Type I and Type II Pores of Arabidopsis Nodulin 26-Intrinsic Proteins"

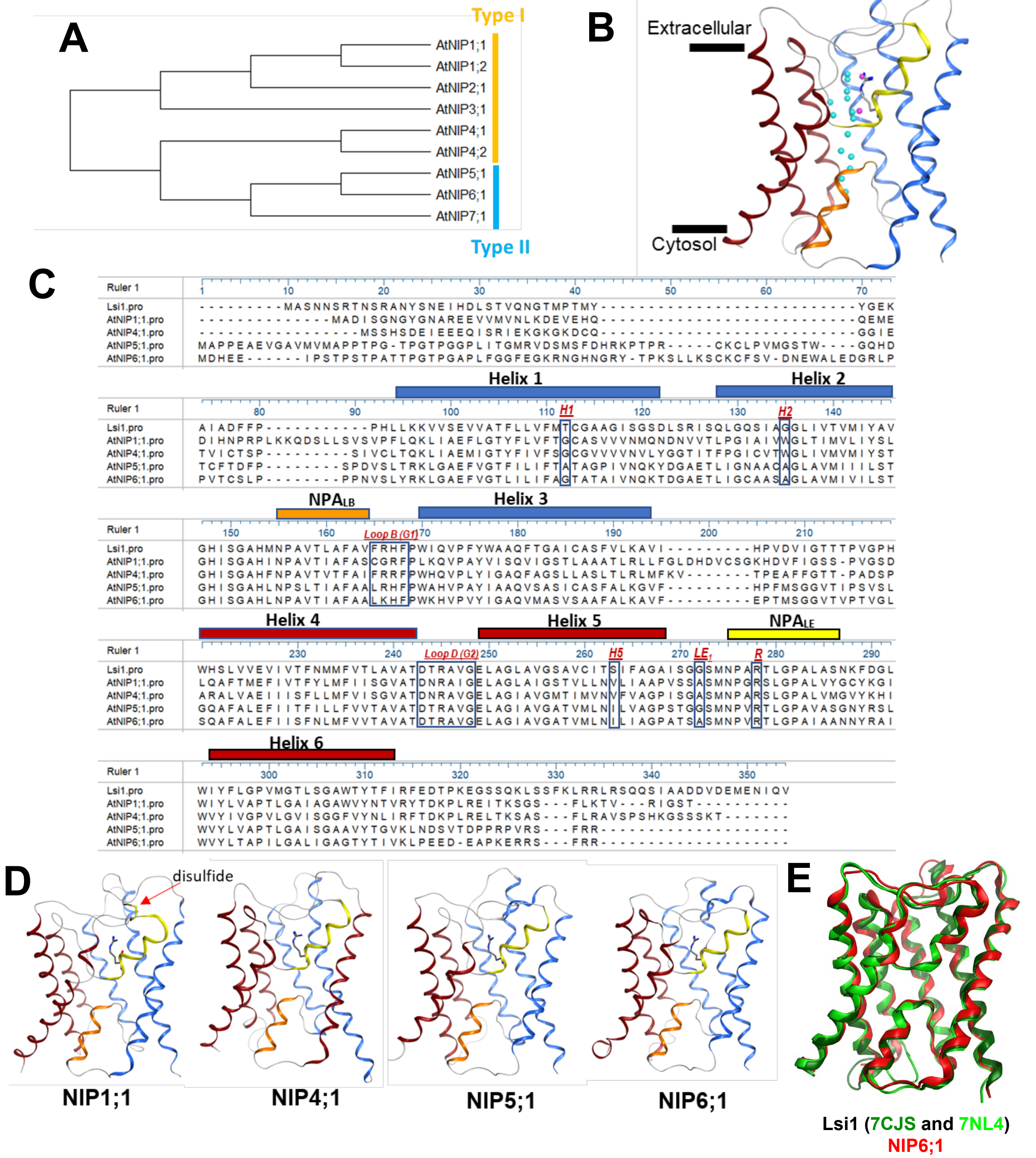


**Figure S1 Alignment and Alpha-fold modeling of Arabidopsis NIP I and II proteins. A.** Phylogenetic tree of the seven *Arabidopsis thaliana* NIP proteins as well as their pore subtype classification. **B.** Sideview of Lsi1 monomer (Saitoh et. al. 2021, PDB 7CJS) with pore water molecules (light blue and pink spheres) and selectivity filter Arg shown. **C.** Alignment of Arabidopsis NIP1;1, NIP2;1, NIP4;1 & NIP6;1 with Oryza sativa NIP2;1 (Lsi1). The five selectivity residues of Lsi1 are denoted by H1, H2, H5, $LE_1$ and R (conserved arg at $LE_2$). The residues responsible for the intracellular gating in the Lsi1 crystal structure (van der Berg et. al. 2021) are denoted by Loop B (G1) and Loop D (G2) boxes. Helices are colored based on the Lsi1 structure (panel B). **D.** Ribbon structures of the indicated alpha-fold model structures with the same color scheme and the position of the ar/R arg side chain highlighted. The predicted position of a disulfide bond between loop C and B in NIP1;1 is indicated. **E.** Superimposed structures of NIP6;1 homology model obtained from AlphaFold (red) with the open (dark green, PDB 7CJS) and closed structures (light green, PDB 7NL4) of Lsi1.

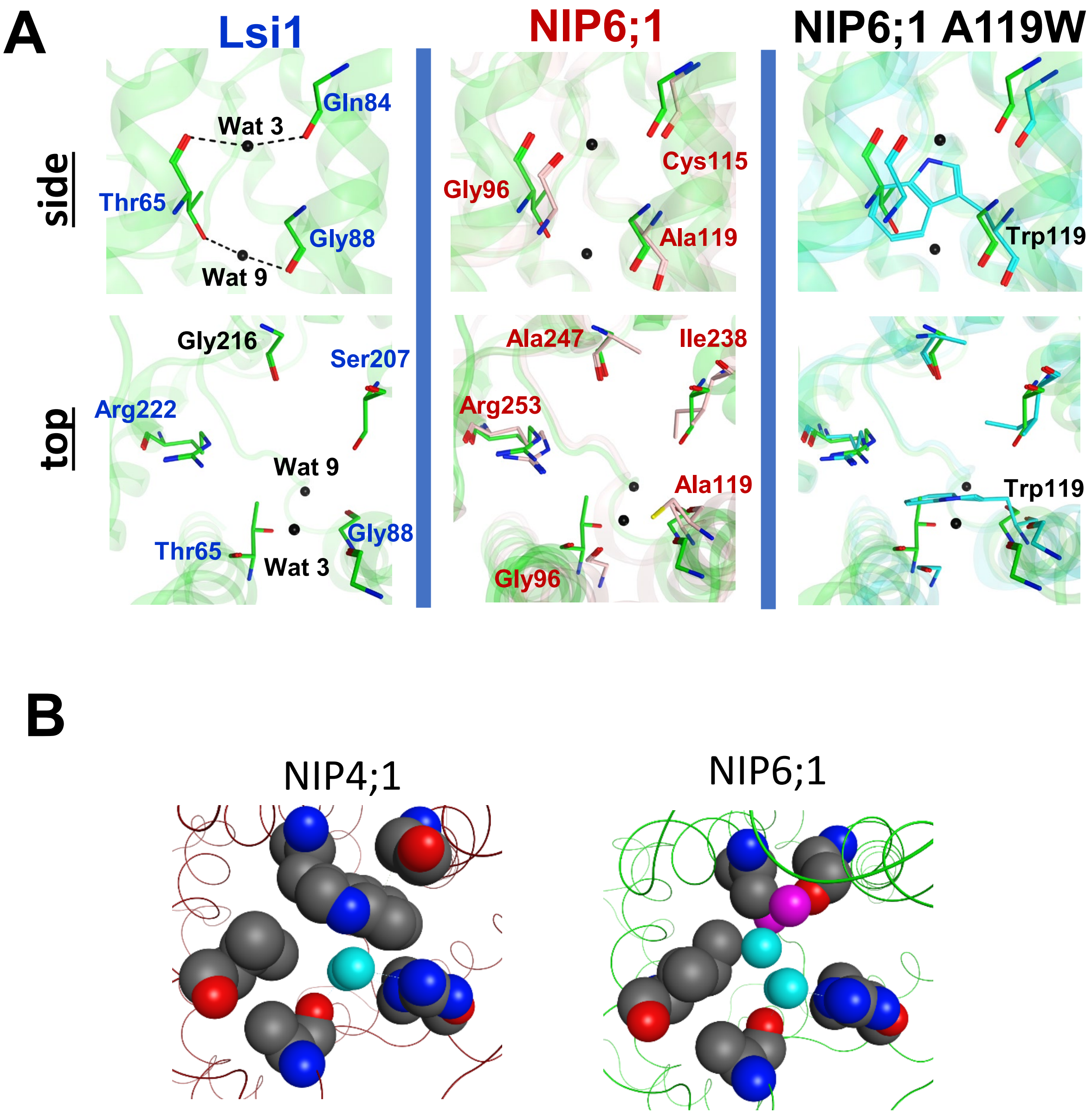


**Figure S2 Effect of H2 tryptophan substitution in NIP I pores. A.** The position of immobile waters 3 and 9 in the Lsi1 structure and the NIP6;1 and NIP6;1 A119W viewed from the membrane plane (upper panel) and extracellular vestibule (lower panel). **B.** Comparison of NIP6;1 (NIP II) and NIP4;1 (NIP1) selectivity filter models (viewed perpendicular to the pore axis from the extracellular vestibule) with waters (aqua and magenta spheres) bound as in Fig. 1C.

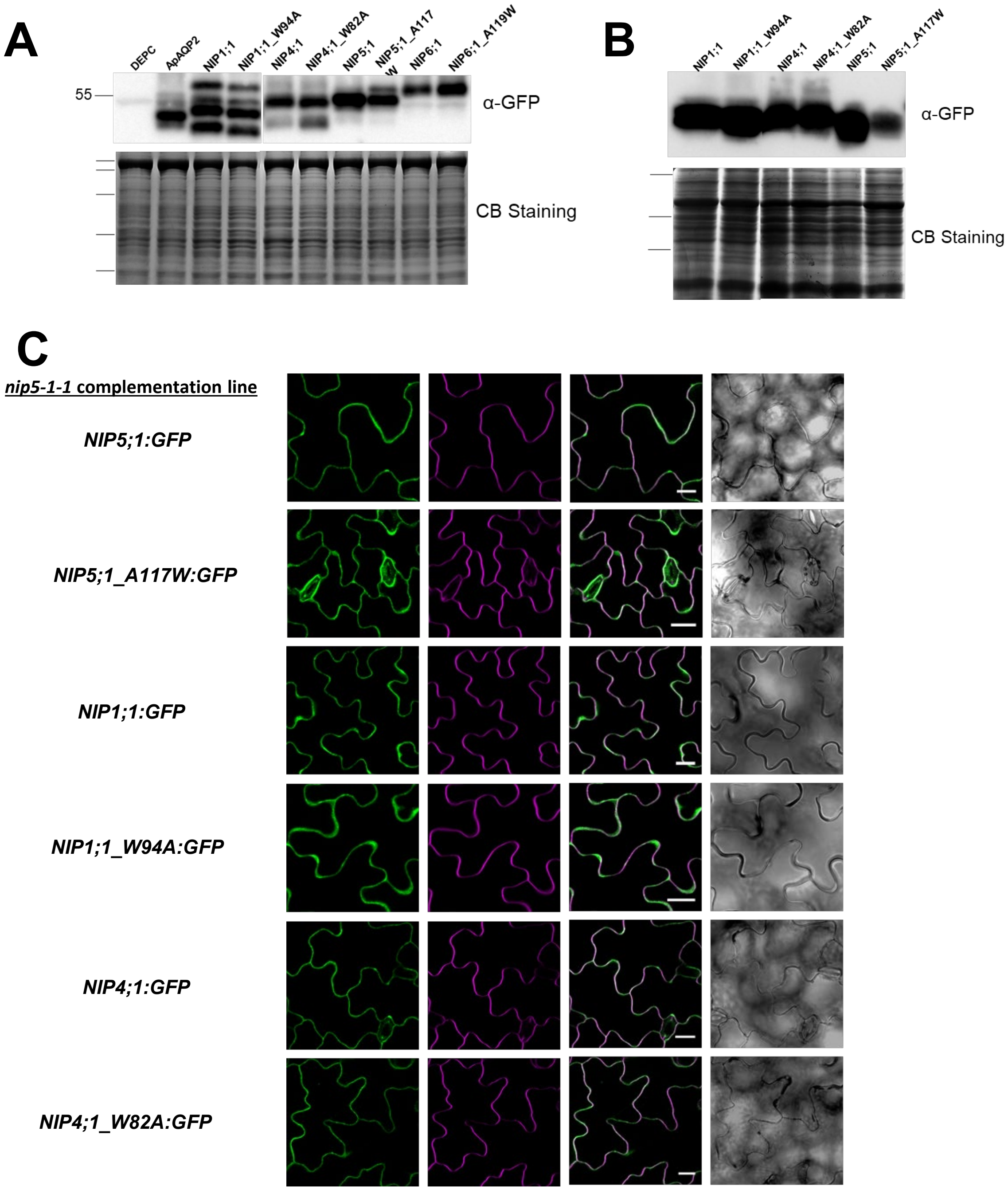


**Figure S3 NIP-GFP expression in *Xenopus oocytes* and *Arabidopsis thaliana* transgenic plants. A.** Anti-GFP Western blot showing NIP-GFP protein in oocytes injected with the indicated cRNAs (upper panel). Bottom panel, Coomassie blue stained loading control gel. DEPC represents negative control oocytes injected with sterile DEPC instead of cRNA. ApAQP2-GFP represents oocytes injected with **B.** Anti-GFP Western blot from extracts of 10 d old Arabidopsis seedlings (upper panel). Bottom panel, Coomassie blue stained loading control gel. **C.** Leaves from seven-day old transgenic *nip5;1-1* Arabidopsis seedlings complemented with the indicated construct were dissected, stained with FM4-64, and were imaged by confocal microscopy. GFP (first panel), FM4-64 (second panel), merged images (third panel) and DIC (fourth panel). Scale bar = 20 μm.

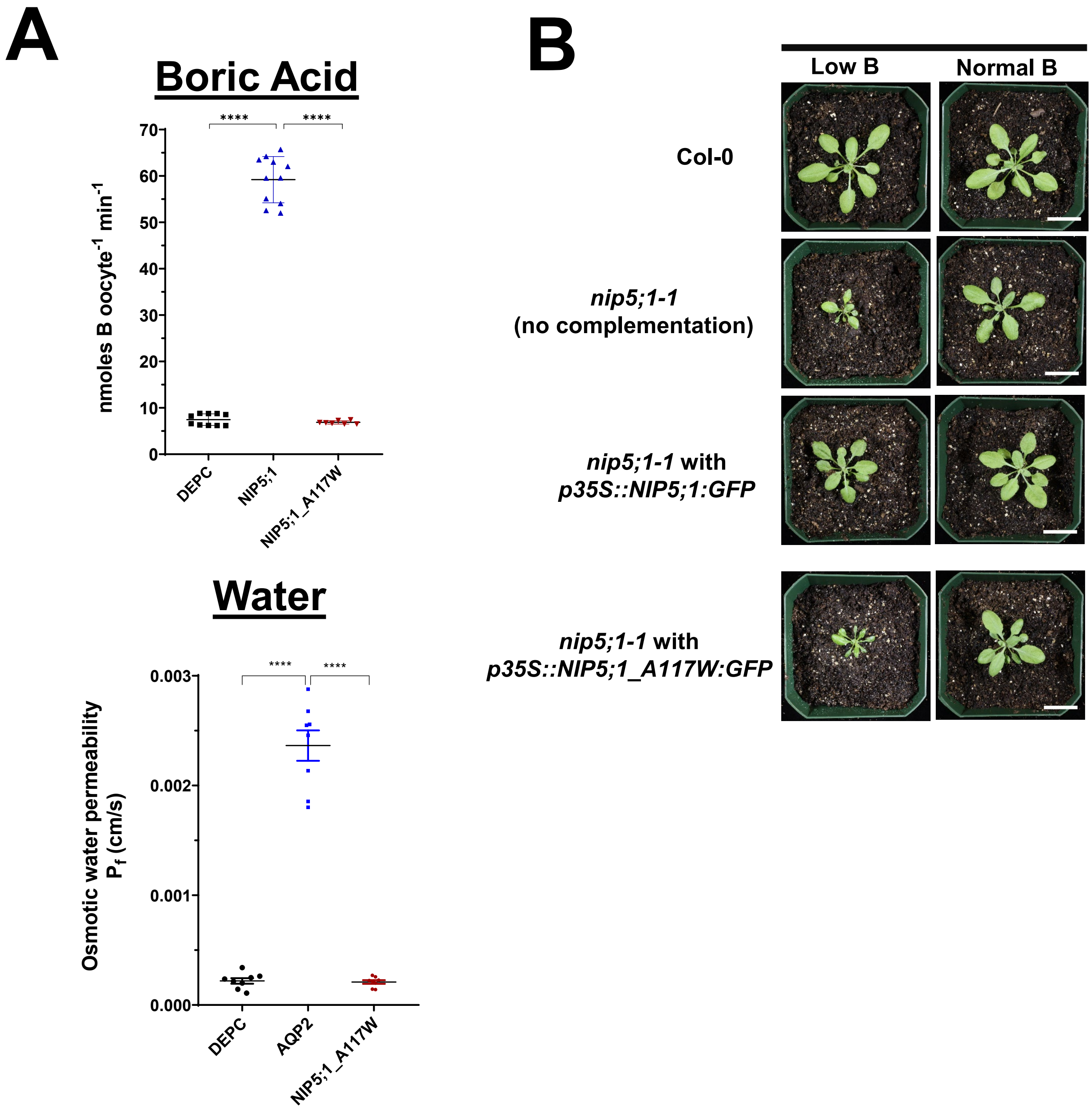


**Figure S4 The H2 mutant protein NIP5;1 A177W produces an inactive protein. A.** Comparison of the boric acid and water permeability of NIP5;1 A117W to positive control (NIP5;1 and the aphid aquaporin ApAQP2, respectively) and negative control (DEPC oocytes. Stastical comparison was done by One Way ANOVA analysis. **B.** Complementation analysis of the *nip5;1-1* B sensitive phenotype with wild type and mutant NIP5;1-GFP constructs**.** Plants represent 29-day old wild type (Col-0), *nip5;1-1* or *nip5;1-1* complemented with the indicated *NIP5;1-GFP* constructs driven by the 35S promotor were grown under low (1 μM) or normal (30 μM) boric acid conditions. Scale bars are 2 cm.

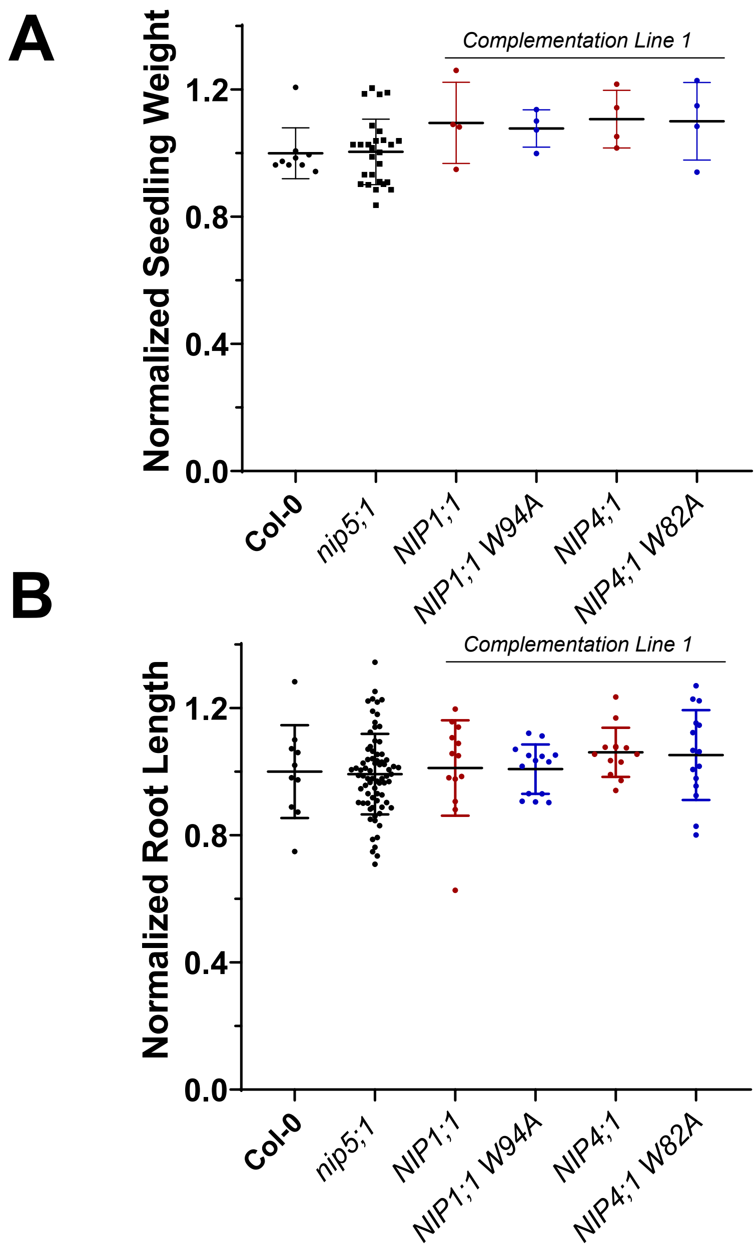


**Figure S5 Comparison of growth of wild type, *nip5;1-1*, and complementation lines under sufficient boric acid conditions.** Seedling fresh weight (A) and primary root lengths (B) of 7-day old seedlings grown under non-limiting boric acid conditions (50 μM) conditions (mean and SD shown as a scatter plot of the data).

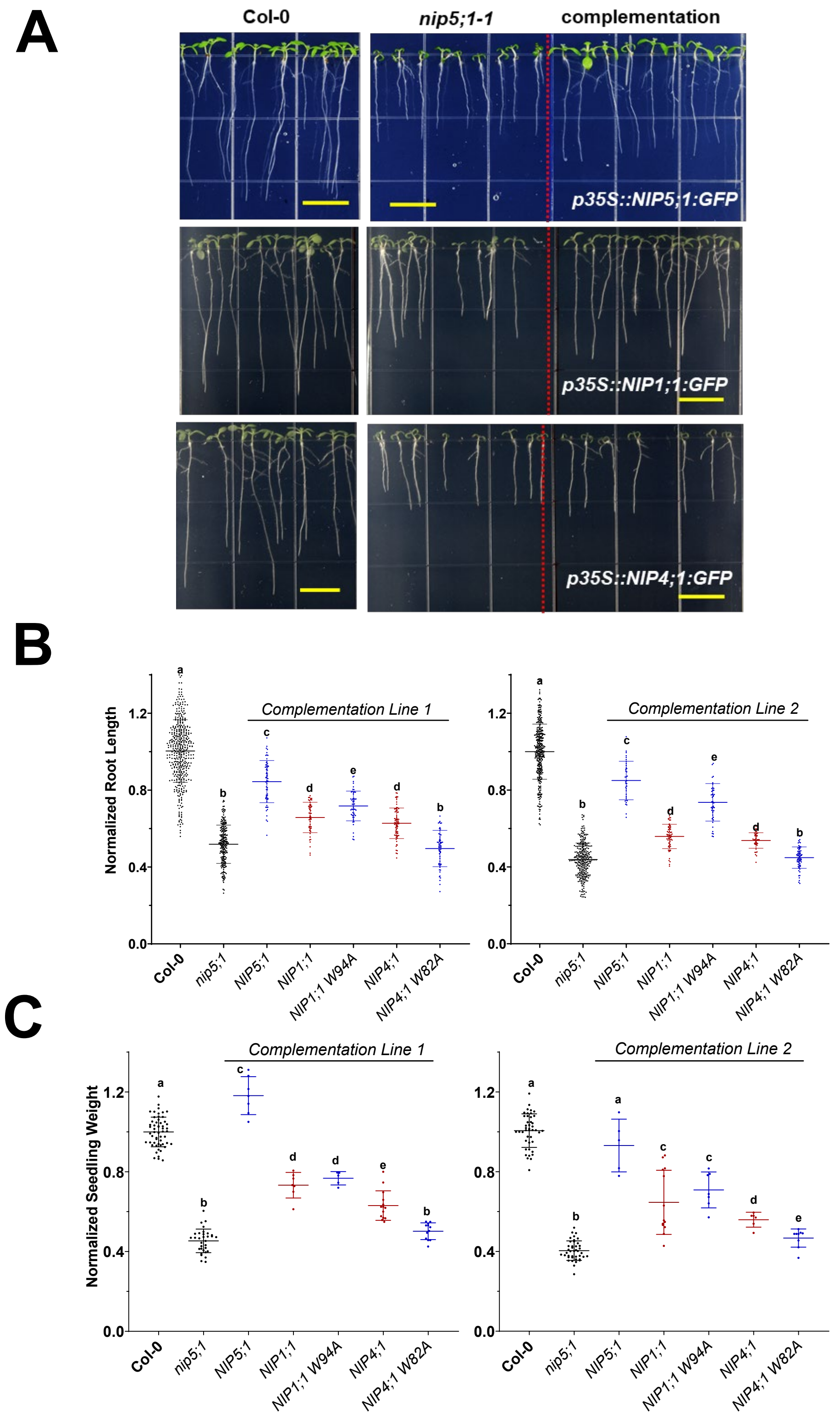


**Figure S6 NIP I protein complementation of the B sensitive phenotype of *nip5;1*. A.** Representative seven-day old Arabidopsis seedlings growth on limiting (1 µM) boric acid media. **Col-0**, wild type control; ***nip5;1-1***, *nip5;1-1* plants without complementation; **complementation**, *nip5;1-1* plants complemented with the indicated constructs (complementation line 1). **B.** Comparison of the primary root lengths of seven-day old Arabidopsis plants cultured as in panel A as measured in ImageJ software and represented as a scatter plot (mean and SD). Each datapoint represents a single seedling. The results of experiments from two representative complementation lines are shown. **C.** Seedlings grown as in panel A were collected in pools and fresh weights were recorded. The mean weight of the Col-0 control value was used to normalize the seedling weight values (each data point is a single pool). Statistical significance ($P<0.05$) was determined by One-way ANOVA analysis with different letters indicating statistically significant difference.

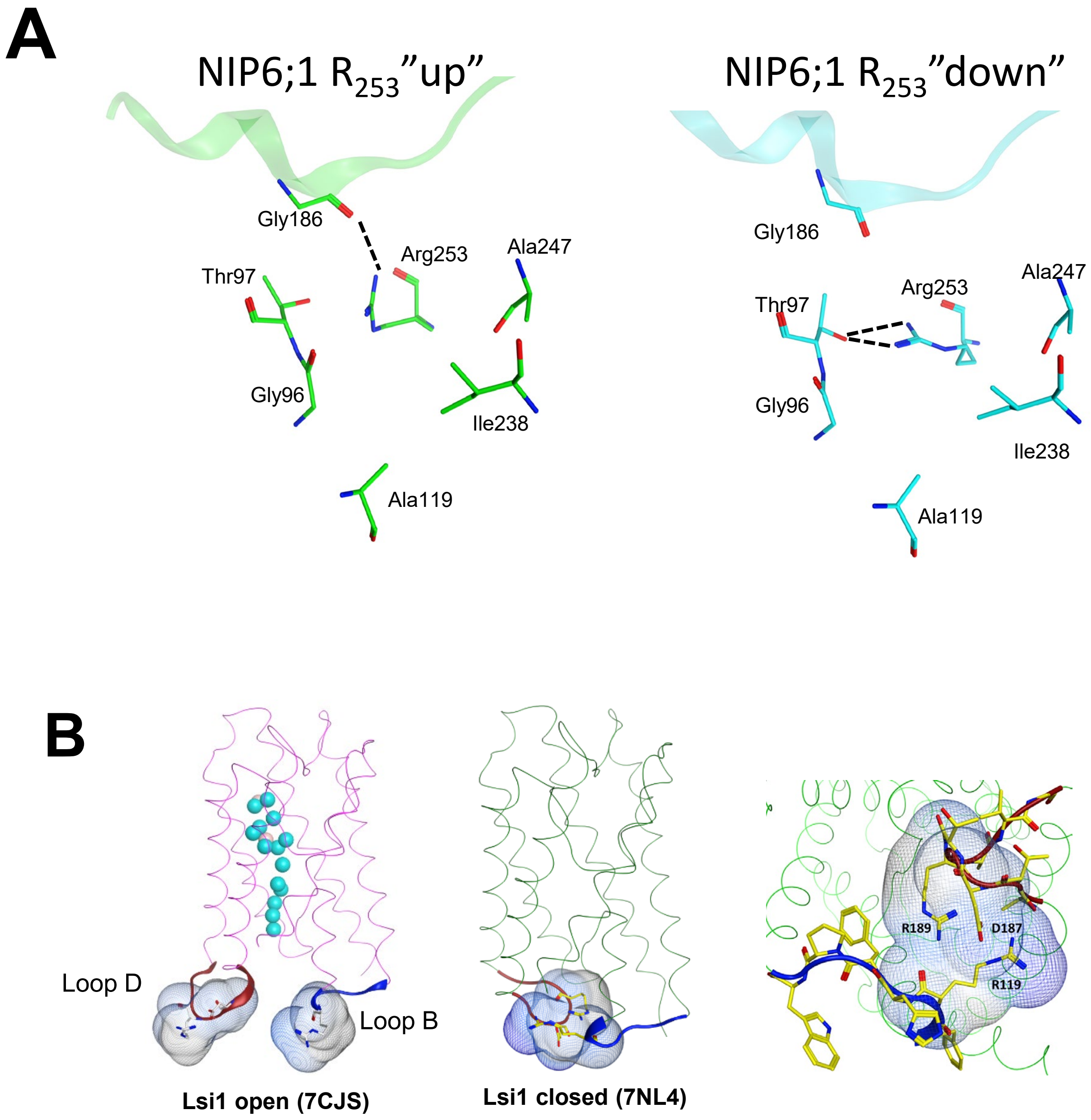


**Figure S7 Potential NIP6;1 and Lsi1 gating conformations. A.** NIP6;1 selectivity filter Arginine occupies two conformational states during MD simulations. Up and down orientation of the selectivity filter arginine (R253) in NIP6;1 homology models identified during MD simulations. Gly 186 is a conserved glycine residue found in the central C-loop of NIP proteins and other aquaporins that stabilizes the arginine in an "up" configuration that maximizes pore diameter. The down conformation is stabilized by a conserved threonine in NIP II proteins within transmembrane alpha helix 1 (see Fig. S1C). **B.** The Lsi1 silicic acid channel structure shown in the open conformation with 16 pore water molecules (Saitoh et al., 2021, pdb 7CJS) and in the closed conformation obtained with $Cd^{2+}$ (van den Berg et al., 2021 pdb 7NL4). The positions of the cytosolic loop D and B regions (see Fig. S1 for reference) are shown.
